# MicroRNA regulation of CTP synthase and cytoophidium in *Drosophila melanogaster*

**DOI:** 10.1101/393181

**Authors:** Najat Dzaki, Woo Wai Kan, Ghows Azzam

**Author notes:** Equal contribution (Najat Dzaki, Woo Wai Kan). Corresponding author (G.A).

## Abstract

CTPsyn is a crucial metabolic enzyme which synthesizes CTP molecules through the *de novo* or salvage pathway. It has the extraordinary ability to compartmentalize into filaments termed cytoophidia. Although this property is retained amongst orthologues, and cytoophidia are therefore found across kingdoms, the mechanisms behind their formation remain unknown. Micro-RNAs (miRNAs) are single-stranded RNA with length of 20 – 22 nucleotides, capable of exerting mRNA silencing and degradation as a form of regulation. *D. melanogaster* itself has a high total gene count to miRNA gene number ratio, alluding to the possibility that *CTPsyn* may too come under the regulatory effects of these small RNAs. A thorough miRNA overexpression involving 123 UAS-miRNA lines, followed by staining of ovarian cytoophidia *dme-*egg chambers, revealed a small group of candidates which confer either a lengthening or truncating effect on the structure. Prime candidates are identified on the basis of consistency. MiR-975 and miR-1014 are both cytoophidia-elongating, whereas miR-190 and miR-932 are cytoophidia-shortening. Though target prediction shows that miR-975 and miR-932 do indeed have binding sites on *CTPsyn* mRNA, *in vitro* assays instead revealed that none of the four candidates may actually do so. This suggests that the effects asserted by overexpressed miRNAs indirectly reach CTPsyn and its cytoophidia through the actions of middling elements. *In silico* target prediction and qPCR quantification indicated that, at least for miR-932 and miR-1014, these undetermined elements may be players in fat metabolism. This is the first study to thoroughly investigate miRNAs in connection to CTPsyn expression and activity in any species. The findings presented could serve as a basis for further queries into not only the fundamental aspects of the enzyme’s regulation, but may uncover new facets of closely related pathways as well.

## 1.0 Introduction

No biotic life is capable of surviving without carrying out the chemical processes and energy changes known as metabolism. Disruption to metabolic enzymes are known to have adverse effects, including decreased muscular plasticity and higher risk of premature death among the young (Schnurr, Yin, and Scott, 2014), as well as increased chances of developing chronic diseases in later life (Ashrafi, 2007; Dang, 2012; Tuck, 1990). One such metabolic enzyme is CTP Synthase (CTPsyn). Its importance lies in the importance of its product. The enzyme synthesizes the cytidine triphosphate (CTP) nucleotide (Kammen and Hurlbert, 1959; Lieberman, 1956), which is not only one of the building blocks of our DNA and RNA, but is a known energy source and cofactor of other enzymatic reactions (Blackburn *et al.*, 2006). The nucleotide also acts as the seed molecule within the phospholipid and sialoglycoprotein synthesis pathways (Yang, Bruno, and Carman, 1996).

A property which distinguishes CTPsyn from other synthases is its ability to tether into filamentous structures called cytoophidia (Liu, 2010). These are observed in a number of organisms including some bacteria (Ingerson-Mahar *et al.*, 2010), yeast, mice (Noree *et al.*, 2010) and humans (Chen *et al.*, 2011). In *D. melanogaster*, the filaments are found in multiple tissues (Liu, 2010), formed singularly by one of the fly’s three CTPsyn isoforms i.e. Isoform C (CTPsynIsoC) (Azzam and Liu, 2013). Cytoophidia are particularly abundant in its ovaries, easily discernible in both follicle and nurse cells of egg chambers. Ovarian cytoophidia are known to change dynamically in response to microenvironmental factors, both morphologically and numerically. For instance, they are both elongated and more numerous in deteriorating cells (Liu, 2010). Meanwhile, the knockdown of the proto-oncogene *Myc* has been shown to significantly shorten micro-cytoophidia within follicle cells (Aughey, Grice, and Liu, 2016).

Whilst the cross-kingdom conservation of cytoophidia indicates that the structure may play vital roles, what these may be largely remains a mystery. In humans, filamentation increases the catalytic capability of CTPsyn by positioning them into conformations which facilitate their activity (Lynch *et al.*, 2017). However, the opposite is true in *D. melanogaster*, whereby the structure’s formation has instead been shown to act as a means to negatively regulate *dme*CTPsyn activity (Aughey and Liu, 2015). The *dme*cytoophidium is also heterogeneous: immunostaining has indicated that it is made of more than just CTPsyn molecules, though these too are yet to be revealed (Liu, 2011). What, then, are the regulatory elements governing CTPsyn and its associated functions? More importantly, how do they do so?

*D. melanogaster* carries a relatively large proportion of microRNA (miRNA) genes. It is believed that these powerful attenuators of translation could be responsible for the regulation of 90% of its protein-coding genes. Many have been identified in various roles throughout the fly’s development, including cell fate specification (Li *et al.*, 2006), tissue growth and cell survival (Brennecke *et al.*, 2003), as well as aging and neurodegeneration (NLiu *et al.*, 2012). Furthermore, multiple metabolic processes are regulated by these small RNAs. For example, a trinity of fat, steroid, and insulin pathways are reliant on the interplay between *dme-*miR-8 and *dme-*mir-14 (Jin, Kim, and Hyun, 2012; Karres *et al.*, 2007; Varghese and Cohen, 2007).

In our study, we screened 234 miRNA overexpression lines using GAL4 drivers active in ovaries of *D. melanogaster* to examine the effects of overexpression of individual miRNAs towards CTPsyn-cytoophidia. Fifteen were found to either shorten or elongate these filaments, though ultimately only four were found to be consistent enough to warrant further investigation. Whilst it was subsequently found that none of these may elicit *CTPsyn* mRNA directly, *in silico* target prediction followed immediately by prediction-validation by way of qPCR analysis implicated pathways which may be linked to cytoophidia formation. It is hoped that the knowledge provided by this study will go on to aid future breakthroughs concerning CTPsyn and cytoophidia regulation.

## 2.0 Materials and Methods

### 2.1 Fly rearing and maintenance

All stocks were raised at 25°C on standard cornmeal-based food, modified to fit the local climate and availability of ingredients. Flies are transferred onto fresh food every three weeks. Weak lines are propagated with the addition of wet yeast.

### 2.2 Selection of driver and miRNA lines

Oregon-R are used as wild-type controls in all experiments unless indicated otherwise. Drivers were chosen on the basis of expression patterns, strength, and relatability to the objectives of an experiment. Expression localization is determined through crosses of each individual driver-GAL4 line to a *w[*] ; UAS-IVS-mCD8::GFP* line. A list of stocks used in this thesis are listed in Supplementary Table S1.

### 2.3 miRNA overexpression screen

To generate miRNA-overexpressing flies, lines bearing selected driver-GAL4 constructs are crossed to UAS-bearing lines carrying an extra copy of either a particular or a cluster of pre-miRNA sequences. Follicle cell driver-GAL4 lines #108022 (*P{w[+mW.hs]=GawB}T155*) and #107204 (*C323a-GawB(GAL4), w[1118]*) are crossed to all available miRNA lines. A ubiquitous driver line #107727 (*y[1] w[*];Act5C-GAL4/ CyO*) are crossed only to lines whereby the overexpression of the miRNA does not cause death before eclosion could occur (Schertel, Rutishauser, Förstemann, and Basler, 2012). Line #107748 (*y[1] w[*]; nos-GAL4*) is crossed to all miRNA lines whereby the UAS-construct was inserted with the *p* plasmid variant, which enables germline-cell expression. All crosses are made on food supplicated with wet yeast, with double the number of females to males, and kept at 28°C for the whole duration of mating and growth. Progenies are transferred onto wet yeast two days after eclosion to encourage ovary engorgement.

### 2.4 Immunofluorescence

Females are isolated after a maximum of 48 hours on wet yeast. Dissection is conducted in lab-standard PBT (1X PBS with 0.2% Triton-X). Ovaries are fixed with 4% paraformaldehyde (PFA) for 10 minutes, then washed thrice with PBT at room temperature for 20 minutes per cycle. PBT is removed completely before blocking with PBTB (PBT with 5% normal goat serum) for a minimum of one hour. Nutation with primary antibody lasts overnight at room temperature (1:200 of rabbit anti-*dme-* CTPsyn (code #SY4468), courtesy of Liu lab). Ovaries are washed for another three cycles with PBT, then incubated overnight with secondary antibodies (1:2000 of DAPI; 1:500 of DY488 goat anti-rabbit; and 1:500 of DY555 Phalloidin, wherever necessary, and all purchased from Life Technologies, USA). Samples are mounted in secondary antibody and viewed under a laser-scanning confocal microscope (Zeiss LSM710, Oberkochen, Germany).

### 2.5 Plasmid construct production and purification

Two types of constructs are designed to ascertain whether any of the strong candidate miRNAs could elicit *CTPsyn* mRNA directly: (A) one bearing the 3’UTR sequence of *CTPsyn*, and (B) independent overexpression constructs for each miRNA. For A, a 298bp-long sequence immediately downstream of the stop codon of the *CTPsynIsoC* gene was incorporated to the end of the eGFP gene on the selected plasmid backbone (pCaSpeR4; Genbank#: X81645). For B, constructs were incorporated into a plasmid with a ubiquitous promoter (Ac in this case, within pAc360; Genbank#: A13228). Mimic-primary-miRNAs (*m*pri-miRNA) were generated, consisting of each miRNA-gene’s premature sequence along with ∼100bp nucleotides flanking it either way. All were introduced into and propagated within DH5α *E. coli* cells (ThermoFisher Scientifc, USA). Primer details are available in Supplementary Table S2. Plasmids were purified prior to transfection using the Wizard^®^ Plus SV Minipreps DNA Purification System from Promega (Cat. No #A1330, USA).

### 2.6 S2 cell preparation and transfection

S2 cell-variant i.e. S2R+ are acquired as frozen stock from Drosophila Genomics Resource Centre (DGRC) and revived as instructed. Schneider’s Insect medium (Sigma Cat. #S0146) enriched with 10% fetal bovine serum (FBS, Gibco Cat. #10270) and 1% Penicillin-Streptomycin (Invitrogen Cat. #15070-063) is used as culturing media. Cells were maintained at 27°C in an incubator without CO_2_. S2 cells at full confluency in T-75 flasks are resuspended and counted. 1 × 10^5^ cells/ml and 2 × 10^4^ cells/ml were seeded into 6-well and 24-well plates, respectively. Plates are returned to the growth incubator for 20 to 24 hours prior to transfection. Effectene^®^ (Qiagen Cat. #301425) is used exclusively as the transfection reagent in ratios recommended within manufacturer’s instructions. Each instance of plasmid introduction is followed by an incubation period of 60 hours.

### 2.7 Flow cytometry

Cells doubly transfected with pTub-eGFP-3’UTR-IsoC and pAc-360-miR*Xov* were fixed with fresh 4% PFA in sterile PBS for 15 minutes at room temperature. Successive steps were as described elsewhere (UToronto, 2017). Signal normalizing controls include (a) empty pTub-eGFP backbone, and (b) pTub-eGFP-3 ′ UTR-IsoC doubly transfected with empty pAc-360 backbone, and (c) S2R+ cells supplicated with Enhancer^®^/Effectene^®^ complex, but without any plasmid DNA. Statistical significance in signal intensity differences were determined by simple Student T-test.

### 2.8 Candidate miRNA target prediction

*In silico* target prediction of candidate miRNAs were conducted through TargetScanFly6 (Kheradpour *et al.*, 2007), with a version last updated in 2015. The top three non-hypothetical proteins listed are considered top hits; however, if a gene within sight is known to be directly related to cellular localization or biogenesis *and* is highly expressed in reproductive tissue i.e. ovaries or testes, it is included over another gene which show neither of those characteristics, despite being a more highly ranked target. Gene ontology details and RNA-seq data were retrieved from publicly available information on FlyBase (Gelbart *et al.*, 1997; Gelbart and Emmert, 2013). Target mRNA-specific primer pairs were designed on the Primer3 (http://bioinfo.ut.ee/primer3-0.4.0/) platform or through NCBI’s own primer-pick function. These are named accordingly, with the acronym of its candidate miRNA preceding the gene, e.g. *m932-geneX*, and are viewable in Supplementary Table S3. Another function of TargetScanFly i.e. ORF was used to corroborate predictions, by introducing the 3′UTR sequence of CTPsyn (CG6854) into the software.

## 3.0 Results

### 3.1 Cytoophidia-affecting miRNAs

In order to look for miRNAs that are affecting CTPsyn, we carried out an overexpression screen of miRNAs and look for the changes in cytoophidium as an indicator. Crosses were made between a virgin driver-GAL4 female and a UAS-*miR-X* male; *X* represents any miRNA. Each cross was repeated at least thrice. Reciprocals were made at random, to assess whether outcomes were sex-influenced. A total of 123 miRNAs borne by 234 fly lines were screened (listed in Supplementary Table S1). Traits of note were primarily cytoophidia length, numbers, and density or compaction. Affected polarity and the discombobulation of cell shape and size were secondarily considered. Screening was carried out in order, i.e. first using the FCDs *T155-GAL4* and *c323a-GAL4*, followed by the *nos-GAL4* germline-cell driver, and lastly the ubiquitous *Act5C-GAL4*. The reference phenotypes utilized as controls for all crosses were driver-dependent i.e. based on the morphology of cytoophidia in *driver-GAL4>Oregon-R* flies. It is worth noting that certain drivers do appear to have minor cytoophidia-altering effects, even without the overexpression of miRNAs (Figure 1).

**Figure 1:**
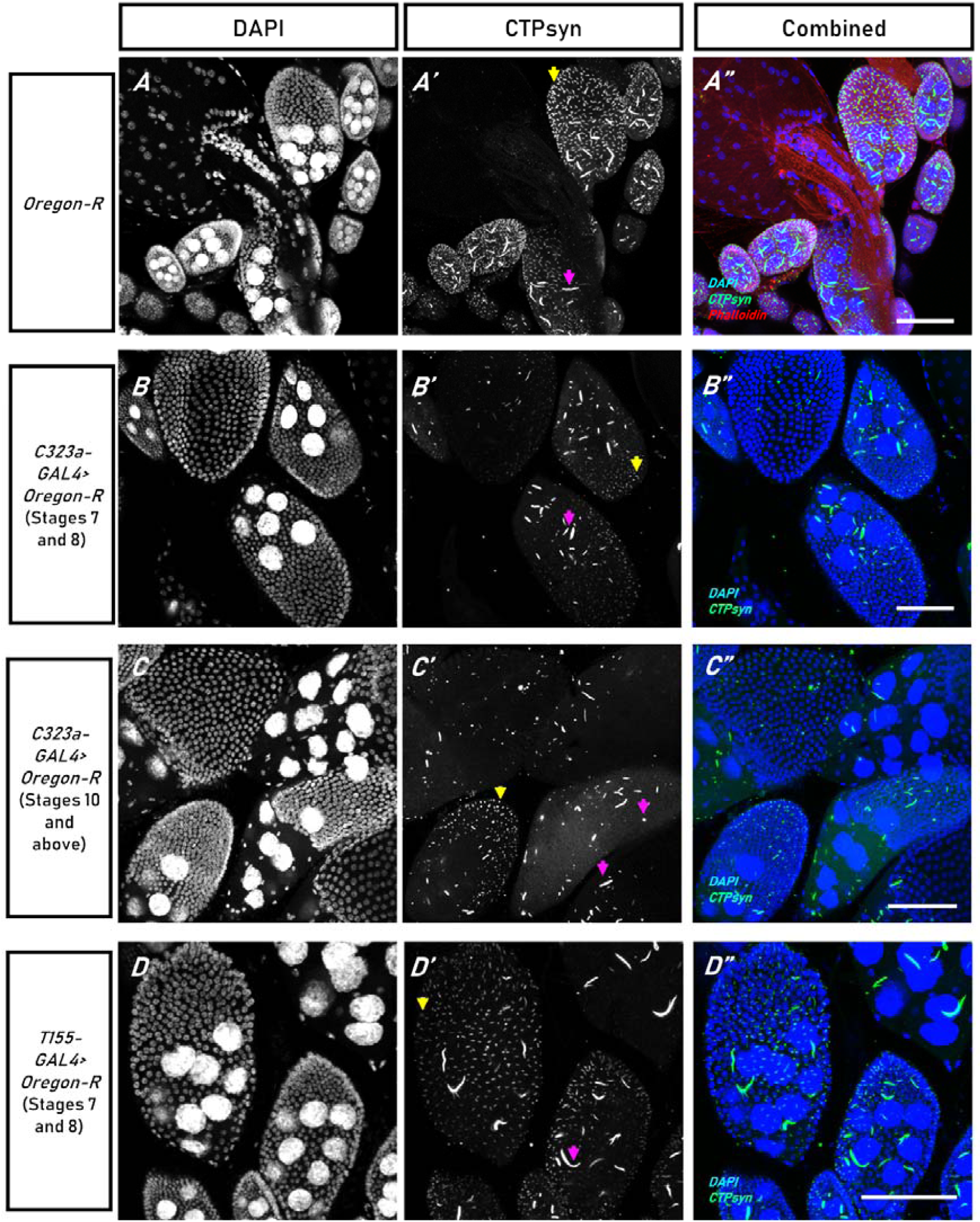
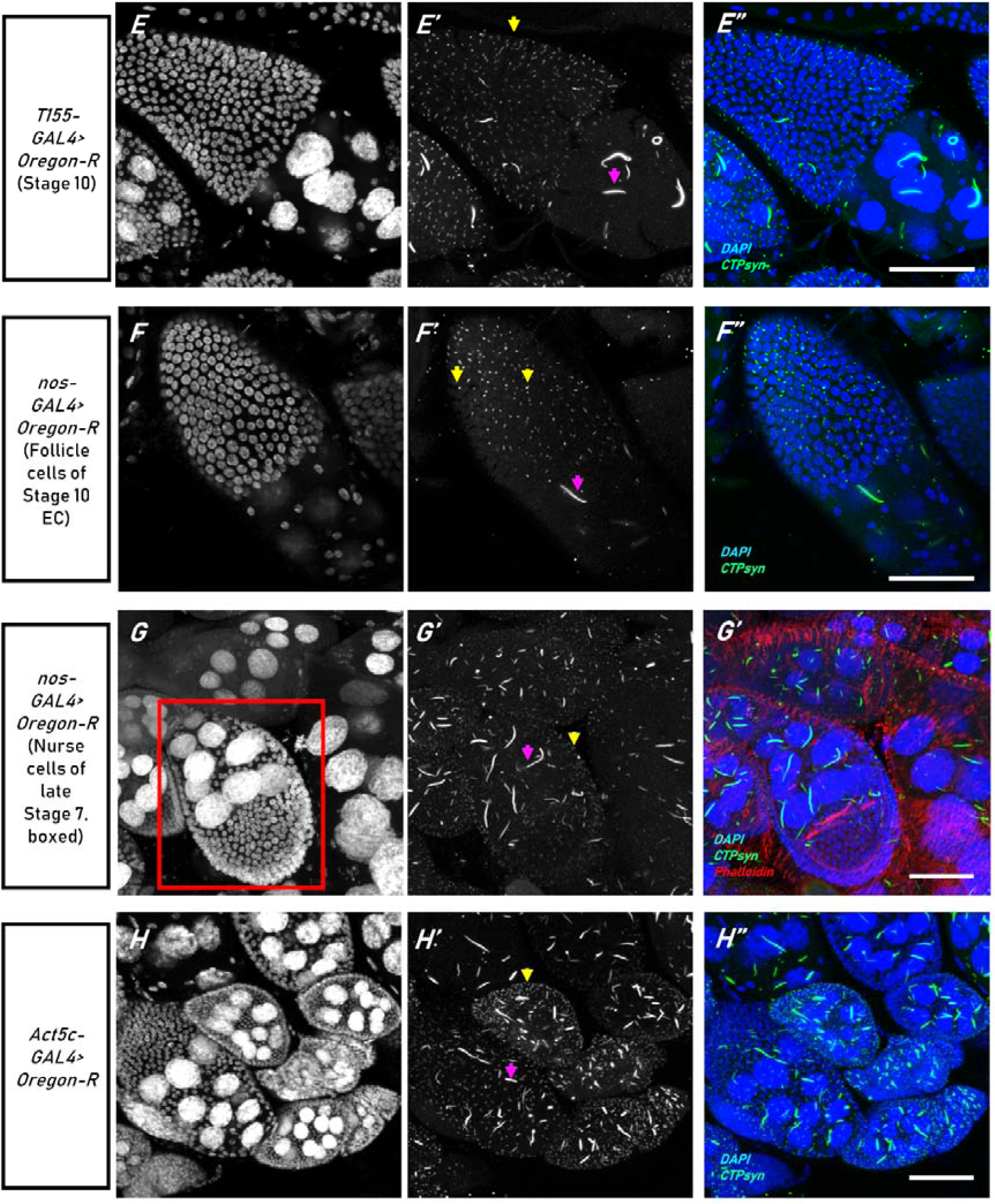
Driver-GAL4 specific reference phenotypes. Genotypes of ovaries displayed are as indicated within side panels; top panels indicate immunostained proteins. CTPsyn form cytoophidia. Here, macro-cytoophidia are indicated by purple arrows, and micro-cytoophidia are indicated by yellow arrows. In B and C, where truncation of the structure has occurred, they may appear as punctuates instead of filaments. All scale bars represent 40μm. (A) shows cytoophidia as they appear in wild-type Oregon-R, presumed to be free of introduced constructs; (B and C) The follicle cell driver (FCD) *C323a-GAL4*, when crossed to Oregon-R, shortened cytoophidia in both follicle and nurse cells, and in egg chambers of various stages. (D and E) *T155-GAL4*, an alternative FCD, also truncates cytoophidia, but only in nurse cells. (F and G) Cytoophidia within germline-specific driver *nos-GAL4* expressing ovaries are unaffected. (H) Cytoophidia in ubiquitous *Act5c*-*GAL4* expressing ovaries are slightly shorter than average.

In the primary stage of screening (with FCDs), a candidate miRNA is shortlisted if its overexpression was observed to cause changes to any of the mentioned traits. If the candidate showed replicability i.e. same phenotype was observed in the next bioreplicates of the cross, it was retained for secondary screening (with *nos* or *Act5c* GAL4-drivers). Some miRNAs were represented by more than one *UAS-miRX* lines i.e. ‘cousin-constructs’. They differ by chromosomal location and point of insertion within the *D. melanogaster* genome. In such cases, the miRNA’s overexpression by the cousin-construct should also produce a similar outcome, although leniency was given in the way of phenotypic strength. Extremity of phenotypic manifestations and reproducibility under the varied array of drivers utilized allowed us to rank cytoophidia-elongating (CytEl) and cytoophidia-shortening (CytSh) miRNAs for the purpose of downstream prioritization. The full list of candidates from all three stages of screening is represented as Table 1. miRNAs marked with an asterisk (*) were eventually shortlisted as prime candidates.

**Table 1:** Summary of expression of miRNA candidates and its characteristics under different drivers, and within different constructs. The candidates for each category are arranged in order of priority. miRNAs marked with an asterisk (*) are further shortlisted as prime candidates.

| Effect | *dme-miR* | with *FCD-GAL4* | with *nos-GAL4* | with *Act5c-GAL4* | Redundancy i.e. performance across all available lines |
| --- | --- | --- | --- | --- | --- |
| **Elongating (CytEI)** | **975\*** | Yes, highly consistent and strong | NA | Yes, high penetrance i.e. >75% of egg chambers | Nine lines carry miR-975. Elongates best on its own (#60663, #3016 and #3026)) or in lines with miR-976/977 (#116509, #60634 and 60635). Attenuated when in a line together with miR-4966 (#41223) |
|  | **1014\*** | Yes, highly consistent and strong | NA | Yes, high penetrance i.e. >75% of egg chambers | Three lines carry miR-1014. Elongates with consistency, but strongest with one line in particular (#3071). Cause higher numbers of cell death |
|  | **276a** | Yes, but not consistent | Yes, but not obvious from control | Yes, but lower penetrance i.e. <40% of egg chambers | Five lines carry miR-276a. Rather iffy with *FCD* but lengthening effects considerably improved under *nos-GAL4*. Low penetrance in egg chambers under *Act5c* |
|  | **10** | Yes, but not consistent | NA | Yes, but low penetrance i.e. <25% egg chambers | Two lines carry miR-969. Most obvious effect with #3108, i.e. non-CyO line., but with a TM3 balancer chromosome. Low penetrance often coincides with low survival i.e. confer a certain degree of lethality under *Act5c* |
|  | **988** | Yes, but not consistent | NA | Yes | Two lines carry miR-988. Neither assert greater effects than the other. Inconsistent with *FCD* |
|  | **8** | Yes, but not consistent | NA | Yes | Two lines carry miR-988. Neither assert greater effects than the other. Weak with *FCD* |
|  | **969** | Yes, but not consistent | NA | Yes, but not consistently | Three lines carry miR-969. Most obvious effect with #3015, i.e. non-CyO line |
|  | **193** | Yes, but not consistent | Yes | Yes, medium low penetrance at ~35% | Three lines carry miR-193, all UASp. Under *FCD*, very little lengthening was discernible; under *nos*, the elongation of cytoophidia became obvious. However, penetrance was unsatisfactory with *Act5c*, and elongation is typically observed alongside heightened lethality |
|  | **1001** | Yes, but barely | NA | Yes, med-penetrance but not all carrier lines produce elongated cytoophidia | One line carries miR-1001. Did not show any out of the ordinary effects under *FCD*, but the phenotype appears with *Act5c*. Lack of a second redundant line disables further steps towards confirmation to be taken |
| **Shortening (CytSh)** | **190\*** | Yes, consistently and highly penetrant | Yes | Yes | Four lines carry miR-190. All four causes shortening to similar degrees. Despite previous reports of its overexpression being lethal, none of the four appear to actually cause a reduction in fitness amongst offspring |
|  | **932\*** | Yes, consistently and highly penetrant | Yes | Yes, but viability is compromised | Three lines carry miR-932. All induce cytoophidia shortening, to similar effects; one line in particular (#3038) reduces viability of F1s considerably |
|  | **87** | Yes in nurse cells but not in follicle cells | Yes; mild (#116431, #3112, #59874) to strong (#116430 and 3119) | Yes, but medium penetrance of ~45% | Five lines carry miR-87. Strongest effects when line #3119 is used in crosses. The others assert noticeably milder effects |
|  | **2a-2, 2a-1, 2b-2** | Yes in both nurse and follicle cells, but poor penetrance i.e. ~20% | Yes, strongly with #3118 and #59849, but medium penetrance i.e. ~45% | Yes, but very low penetrance i.e. ~15%, with compromised viability | Five lines carry this cluster construct. Initially excluded but upon observing their effects in *nos* crosses became a CytSh-candidate. However, penetrance is often unsatisfactory, which shows that the effects of this cluster's overexpression is volatile |
|  | **316** | Yes, but not very strongly in either follicle nor nurse cells | NA | Yes, but low penetrance i.e. ~20% | Two lines carry miR-316. Effects are only mild, with neither construct exceedingly performing the other |
|  | **983-1** | Yes in nurse cells but not as obviously in follicle cells | No | NA (Act5c-lethal) | Two lines carry miR-983-1. In #41217, it is within a cluster with miR-984 and its miR-isoform, 983-2. Shortening only caused when miR-983-1 is singularly overexpressed, i.e. in crosses with line #41194 |

### 3.2 Prime CytEl candidate miRNAs

#### 3.2.1 MiR-975 elongates cytoophidia and forms 32 cell egg chambers

Nine CytEl-miRNAs were identified. Two were considered prime candidates due to their consistency at inducing the lengthening of cytoophidia i.e. miR-975 and miR-1014. Both miRNAs are insect exclusive, and neither have reported human paralogues. The miRNA-975 is carried by 9 lines, in various combinations (Refer to Supplementary Table S1 for full details). On the *Drosophila* genome, its gene is a part of a tri-miRNA cluster, alongside miR-976 and miR-977 (Ryazansky, Gvozdev, and Berezikov, 2011). This cluster is regionally associated with miR-4966 upstream, as well as miR-978 and miR-979 genes downstream, although their expressions are dependent on separate transcription start sites (Figure 2A; (Marco *et al.*, 2013)).

**Figure 2:**
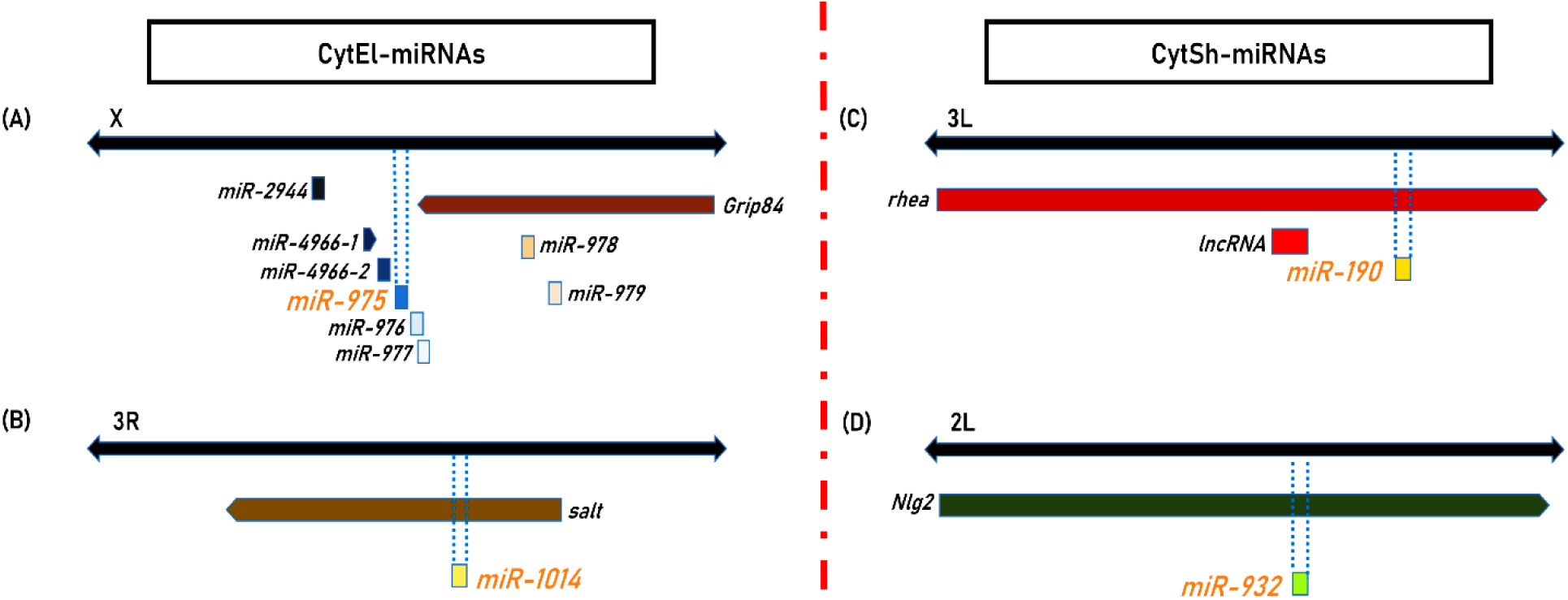
Characteristics of prime candidate miRNA genes. Prime CytEl-miRNAs: (A) miR-975 is encoded in a cluster alongside miR-976 and miR-977, but is additionally flanked within close proximity by miR-4966 and miR-978. (B) miR-1014 is an intronic gene to salt. Prime CytSh-miRNAs: (C) The gene locus of miR-190 is found within the rhea gene. (D) miR-932 is also intronic, found within the ORF of Nlg2.

Throughout the overexpression screen, egg chambers overexpressing miR-975 alone showed significant lengthening of cytoophidia. Whereas elongation was more obvious within nurse cells under FCD-driven expression, the phenotype was obvious in both follicle clearly exhibited by both follicle and nurse cells under *Act5c-GAL4*. Out-of-the-ordinary numbers of micro-cytoophidia did not accompany macro-cytoophidia elongation in either case (Figures 3A and 3B). These observations were repeatable as miR-975 was expressed in a cluster with miR-976 and miR-977, or as a pair with miR-978, though to a lesser degree (Figures 3C and 3D). With miR-975 sharing a construct with miR-4966, it did not cause cytoophidia-lengthening effects, alluding to the possibility that these genes may co-regulate each other’s activity (Figure 3E). However, miR-975 overexpression by the *Act5c-GAL4* driver did not bear cell-death inducing effects. Instead, we have observed the formation of 32 cell egg chamber with a penetration of 2% (n=348) (Figure 3F).

**Figure 3:**
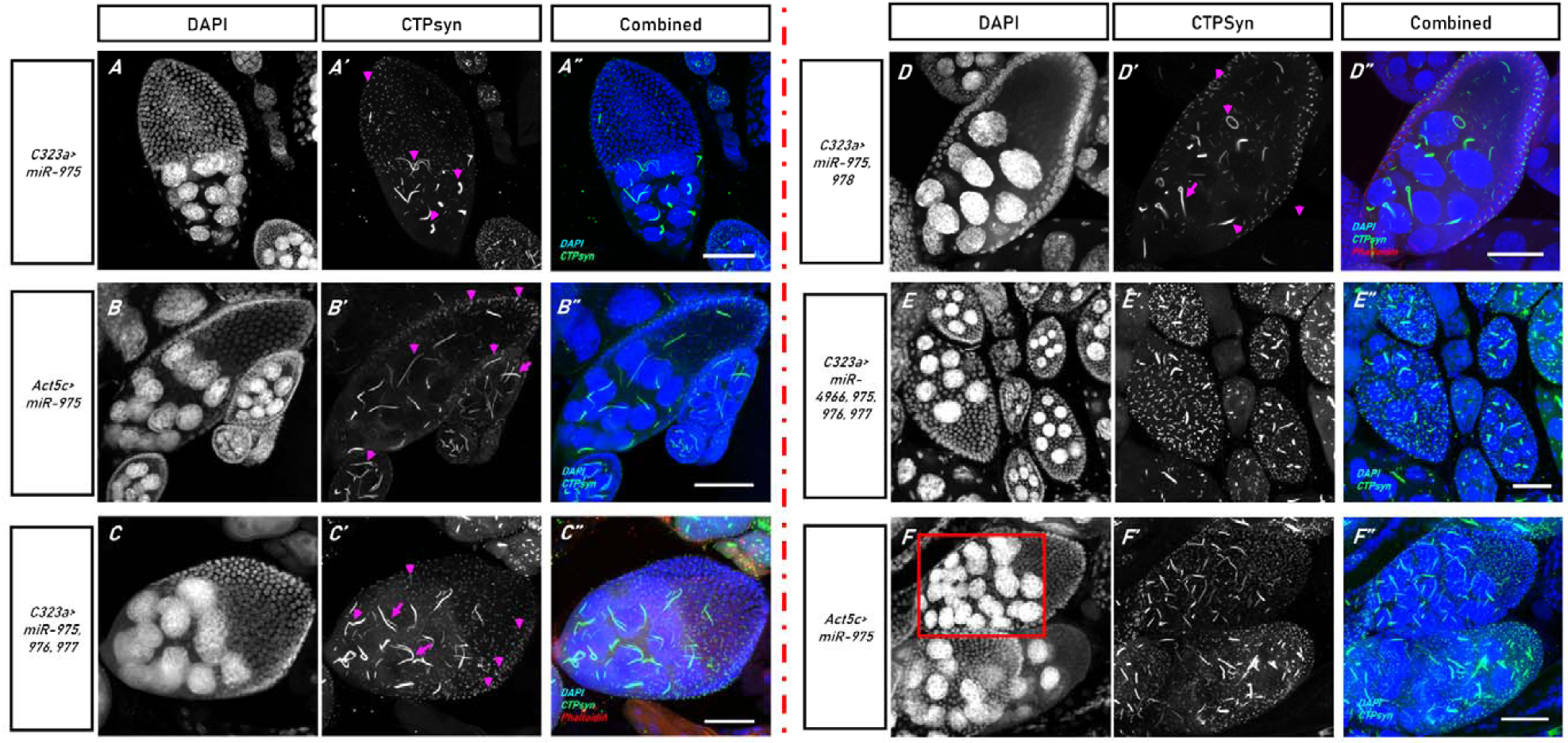
Prime CytEl-miRNA, miR-975, consistently elongates cytoophidia when overexpressed. In (A to D), its effects appeared driver and construct independent. Lengthening, here indicated by purple arrows, is clear in follicle as well as nurse cells, despite all miR-975-containing constructs being UASt-exclusive. In (E), the only instance where miR-975 caused elongation was not palpable was when it was encoded together with miR-4966. However, when miR-975 is alone ubiquitously expressed by Act-GAL4 driver, the doubling of nurse-cell nuclei can occasionally be seen (F, red boxed egg chamber). Genotypes are indicated by side panels, as driver>miR-X. All scale bars represent 40μm.

#### 3.2.2 MiR-1014 elongates cytoophidia and causes cell death

The second CytEl-miRNA identified as a prime candidate is miR-1014, carried by three fly lines. *In vivo*, miR-1014 occupies a locus on chromosome 3, within the intronic regions of the *salt* gene (Figure 2B). Overexpression of miR-1014 consistently induced the elongation of cytoophidia regardless of driver and UAS-construct, though one induced greater effects than the rest (Figure 4A to 4C). Elongation patterns are akin to those described for miR-975. However, excessive miR-1014 levels did appear to negatively impact cell survivability. An estimated 22.35% (n=464) of egg chambers of stage 8 and above were discernably undergoing apoptosis in these ovaries (Figure 4D), a significant increase from natural cell death rates seen in *Act5ct-GAL4>Oregon-R* controls as well as amongst miR-975 overexpressing flies. Nurse cell nuclei doubling phenotypes were not recovered under miR-1014 overexpression.

**Figure 4:**
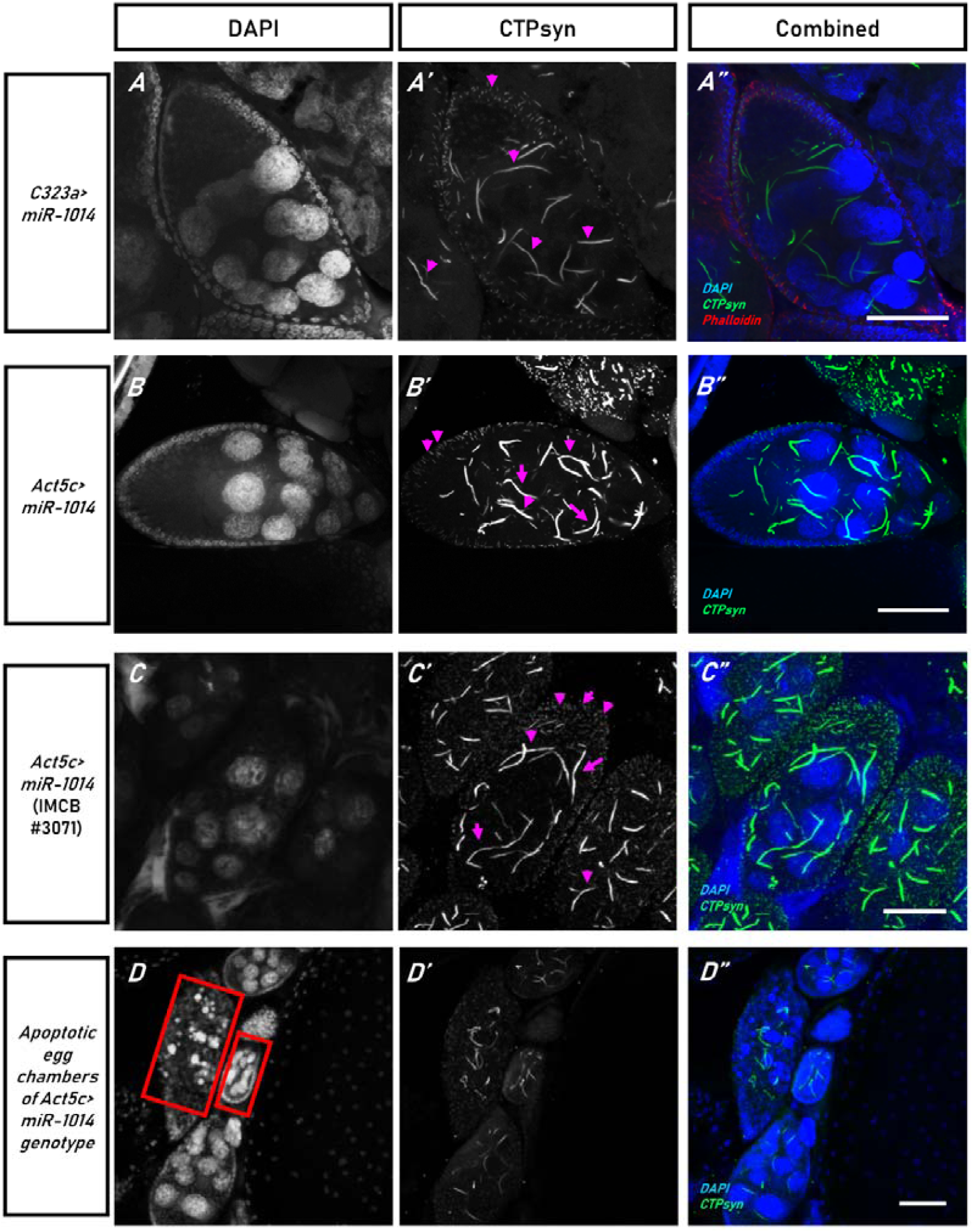
Prime CytEl-miRNA, miR-1014, elongates cytoophidia when overexpressed, but also increases incidence of cell-death. In (A), miR-1014 is driven by an FCD; in (B), it is driven by the ubiquitous driver Act5c. Although all miR-1014 lines are capable of inducing cytoophidia lengthening, this phenotype was most obvious in F1s from crosses with IMCB #3071 in (C), suggesting that severity may be construct-dependent. However, this miRNA’s overexpression significantly increased the frequency of apoptotic egg chambers in comparison to controls and fellow CytEl-miRNA, miR-975 (red-boxed in D). Purple arrows indicate elongated cytoophidia. Genotypes are indicated by side panels, as driver>miR-1014. All scale bars represent 40μm.

### 3.3 Prime CytSh candidate miRNAs

#### 3.3.1 MiR-190 and miR-932 shorten cytoophidia without significant distinguishing side effects

Six miRNAs were determined to shorten cytoophidia. Two were identified as prime CytSh-miRNAs, namely miR-190 and miR-932. Four UASp-lines encode singularly for *miR-190*: two carry the gene on chromosome 2. For the other two, the gene is inserted into chromosome 3, where the endogenous *dme-miR-190* should also be found (Figure 2C). Its locus is intronic to the *rhea* gene. Overexpression of the miRNA exhibit cytoophidia shortening consistently. The shortening of these filaments is often observed alongside its compaction, and increased numbers of micro-cytoophidia.

Its overexpression is most obvious under *nos-GAL4*, followed by *Act5c-GAL4*. As truncation was already observed in crosses involving FCDs, shortening could only be ascertained by the near disappearance of cytoophidia (Figures 5A, 5C and 5E). Placement of the miRNA-gene within UAS-constructs did not significantly affect outcomes (Figures 5B, 5D and 5F). However, though *Act5c*-driven overexpression of miR-190 was expected to be lethal (Schertel *et al.*, 2012), we often recovered healthy flies which displayed neither signs of compromised fitness nor fertility. More importantly, whereas miR-190 overexpression under FCD showed truncation of both follicle and nurse cell cytoophidia, *nos-GAL4* and *Act5c-GAL4-*driven truncation of macro-cytoophidia within germline nurse cells was not accompanied by the shortening of follicle-cell micro-cytoophidia.

**Figure 5:**
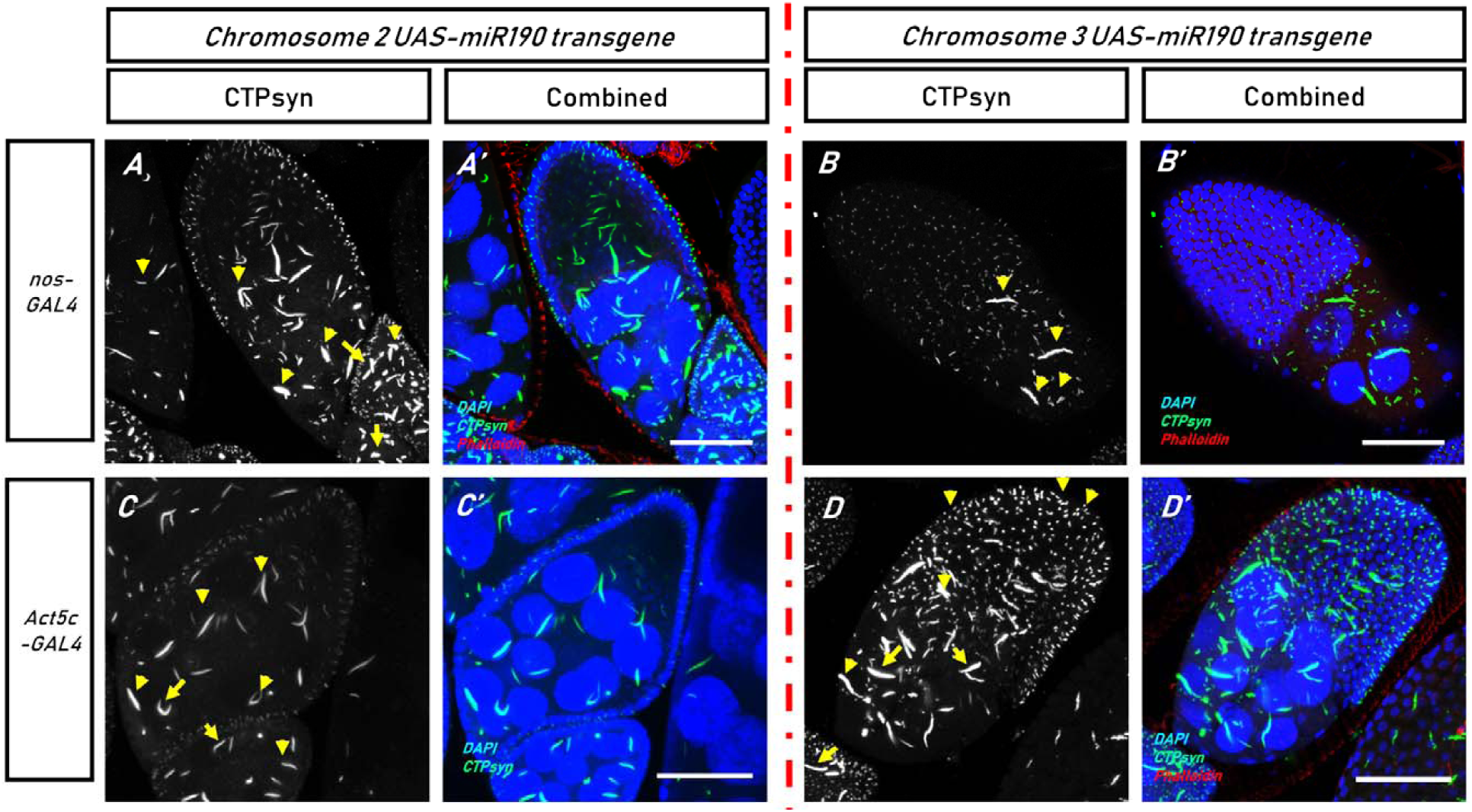
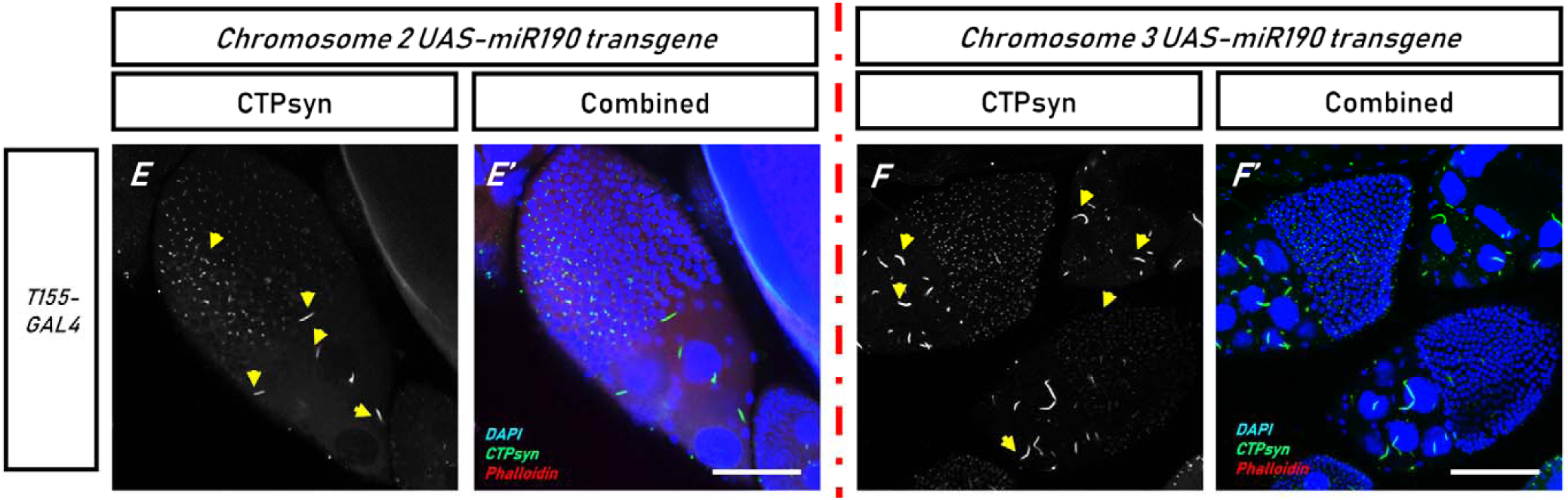
Prime CytSh-miRNA, miR-190, shortens cytoophidia when overexpressed. Cytoophidia truncation appears due to miR-190 overexpression regardless of construct and chromosomal insertion point, but its phenotype is certainly driver-dependent. In (A), (C) and (E), the UAS-miR190 transgene was incorporated into parental chromosome 2; while in (B), (D) and (F), the UAS-miR190 transgene was incorporated into the parental chromosome 3. Parental driver-lines are specified by side panels. Truncated cytoophidia i.e. a positive miR-190-overexpression phenotype is indicated by yellow arrows. All scale bars represent 40μm.

The second prime CtySh-candidate is identified as miR-932. Like miR-975 and miR-1014, the miRNA is also exclusive to arthropods. Three UASp-lines carry this construct. In vivo, the dme-miR-932 gene is intronic to the gene *Nlg2*, found on chromosome 2 (Figure 2D). It induces a severe shortening effect on cytoophidia, as well as an increase in micro-cytoophidia numbers in both nurse cells and the developing oocyte. Shortening of follicle-cell cytoophidia due to overexpression of miR-932 was comparatively more noticeable than that of flies overexpressing miR-190 (Figure 6A to 6D). However, miR-932 carried by the transgenic line IMCB #3038 produces inconsistent phenotypes, where highly-dense, misshapen macro-cytoophidia appeared alongside truncated ones (Figure 6E).

**Figure 6:**
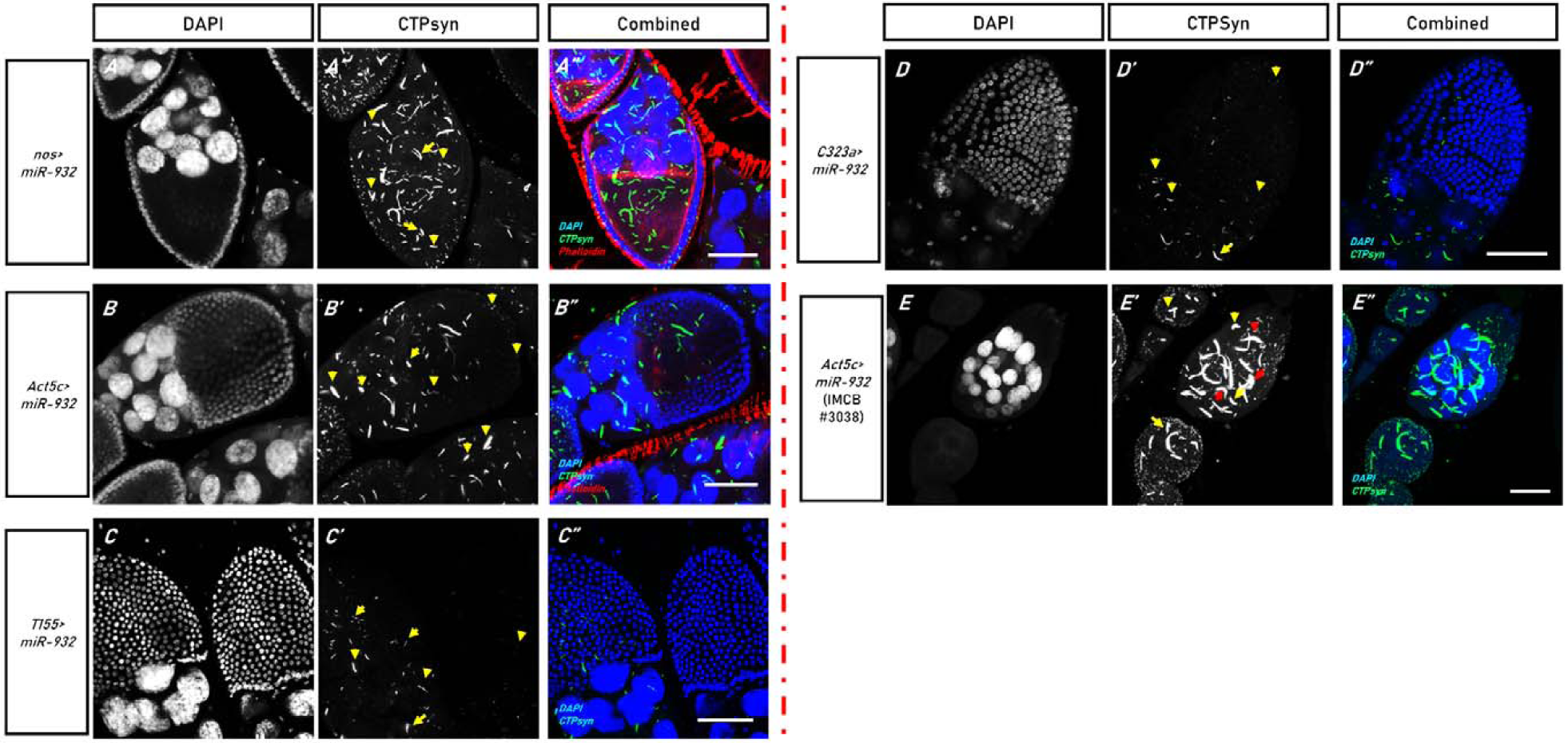
Another CytSh-miRNA, miR-932, shortens cytoophidia when overexpressed. Like miR-190, cytoophidia-shortening phenotype by miR-932 is driver-dependent. In (A) to (D), expression is driven by nos-GAL4, Act5c-GAL4 and each of the FCD-GAL4 drivers, respectively. Effects were considerably consistent; however, in transgenic line IMCB #3038 (UAS-miR932), highly-dense cytoophidia was observed alongside shortened ones in nurse cells (indicated by red arrows in (E)). This indicates that in some instances, different outcomes may be expected due to differences in constructs. Shortened cytoophidia are indicated by yellow arrows. Genotypes are indicated by side panels. All scale bars represent 40μm.

Similar to dme-miR-190, Schertel et al. has also reported that excessive levels of dme-miR-932 would lead to lethality. We found that whilst *Act5c-GAL4>UAS-miR932* flies were viable, their numbers were lower than the F1 produced from the other three prime candidates. This adverse effect on population size was especially true when the *Act5c* driver was crossed to *UASp-miR932* insert in chromosome 2 (Figure 7), with recovered F1 numbers dropping to half of the other two constructs (#3038 vs #3036 vs #116480: 21±4.74 vs 138±11.32 vs 108±9.86).

**Figure 7:**
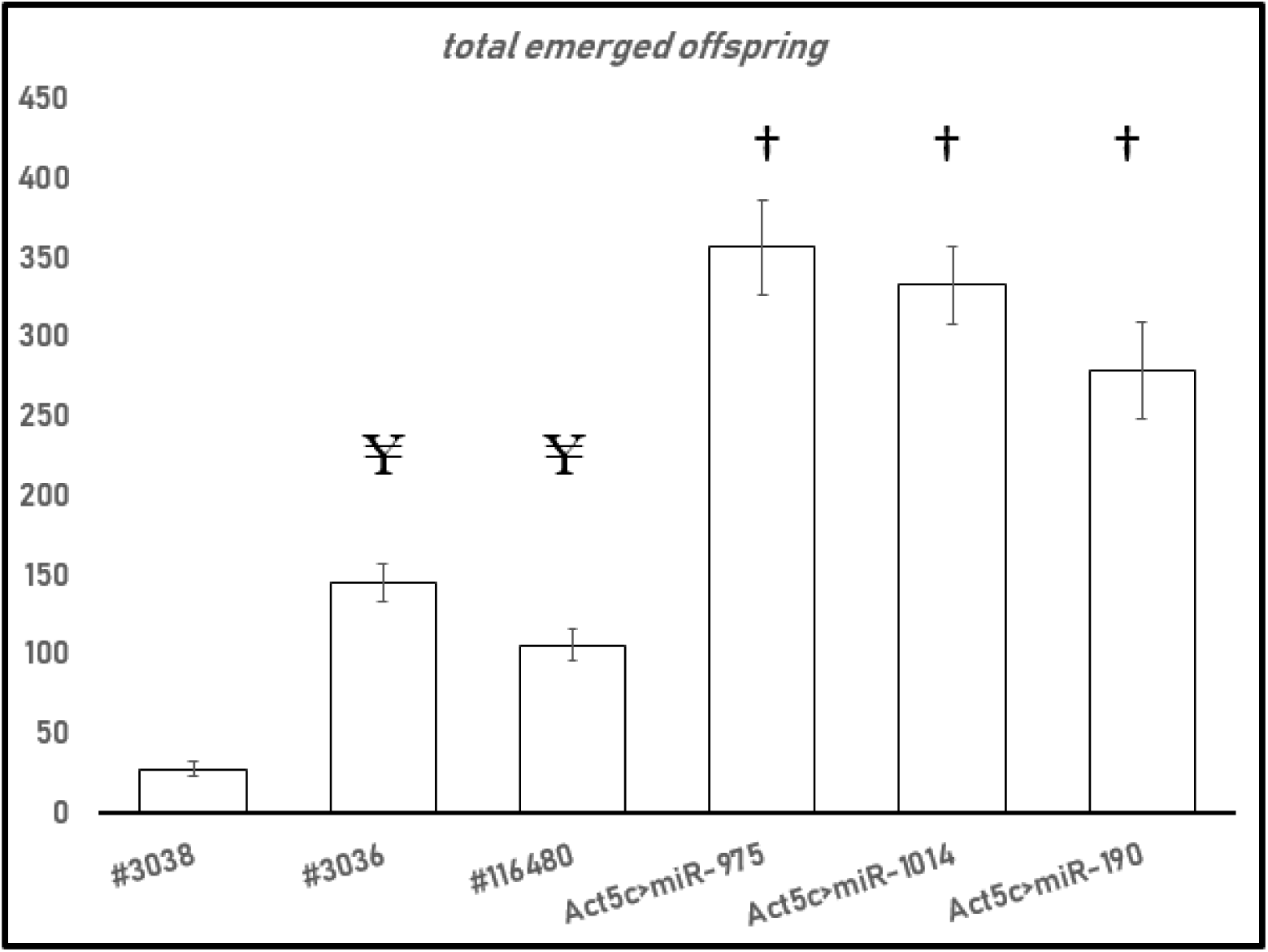
MiR-932 confers a degree of lethality when overexpressed. Schertel et. al (2012) showed that miR-932 to be lethal while being expressed by the ubiquitous driver Act5c. However, our study show that a reduced number of viable offsprings. Total emerged offspring for each of the lines carrying miR-932 construct are compared to other prime candidate miRNAs; and total average numbers as amassed from all of the non-survival affecting constructs. Whilst Student T-Test proves that miR-932 clearly reduces offspring survival in comparison to those miRNAs, it does not exert consistency in its sibling constructs, in which #3038 appears to confer most lethality. ‘†’ symbolizes statistical significance (p<0.05) between miR-932 and other candidates, whereas ‘¥’ symbolizes statistical significance (p<0.05) between #3038 to other mR-932 bearing lines.

### 3.4 Assessing prime candidate miRNA potential for direct regulation of *CTPsynIsoC* mRNA

Activity of overexpression-constructs and its non-lethality to S2R+ cells was confirmed in a preliminary transfection round (see Supplementary Figure S1). These constructs were then co-transfected with a reporter plasmid, which has had the *CTPsynIsoC* mRNA 3′UTR sequence attached to the end of an *eGFP*-gene (Lai, 2002). The proportion of eGFP-positive cells detected through flow cytometry was reduced with the overexpression of all candidate miRNAs. However, T-test confirmed that only the cells transfected with CytEl-miRNA constructs i.e. of miR-975 and miR-1014 (*p<*0.05) showed significant reduction in eGFP signal intensity. Nonetheless, such changes were underwhelming and therefore unconvincing in comparison to controls. 22.97% and 21.55% eGFP-positive cells were detected in miR-975 and miR-1014 overexpressing culture, respectively, compared to the 29.50% eGFP-positive cells in the control culture. Together, these observations suggest that none of the candidates elicit *CTPsynIsoC* mRNA directly. The effects they exert upon the enzyme and its cytoophidia therefore involve mediating parties. Results are summarized in Figure 8.

**Figure 8:**
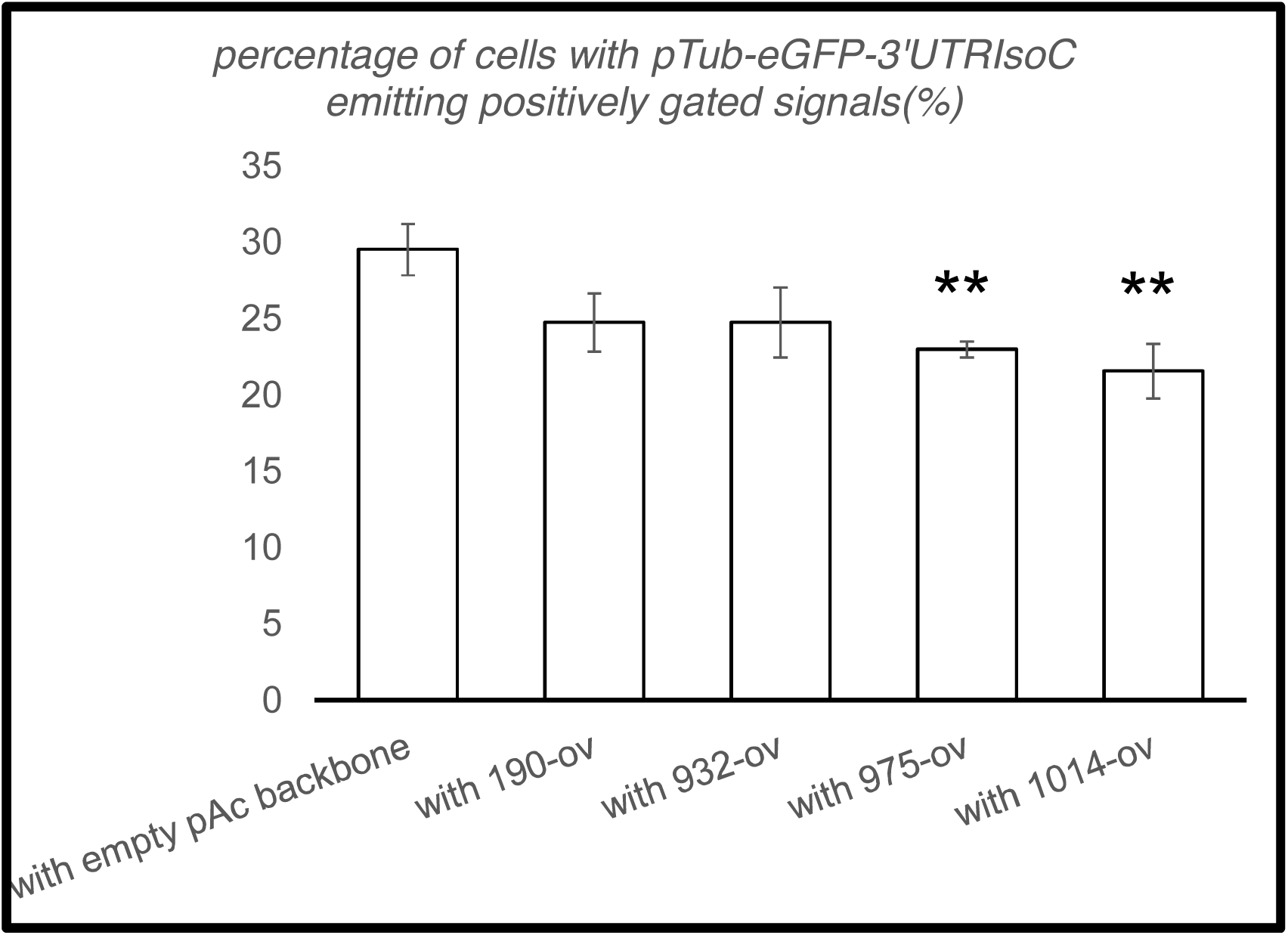
Percentage of S2R+ cells carrying pTub-eGFP-3’UTRIsoC co-transfected with candidate miRNA-overexpression constructs. ‘**’ indicates a significant eGFP signal reduction from control, i.e. cells transfected with an empty pAc-plasmid backbone, without a pri-miRNA sequence.

### 3.5 *In silico* target prediction outcomes

TargetScanFly prediction tool uses complementary-seed-pairing to identify potential targets of any of 148 *dme-*miRNAs within its database (Lewis, Burge, and Bartel, 2005). Only matches to regions in the 3′UTR of *D. melanogaster* mRNAs are considered. Each predicted mRNA target is scored by seed-complementarity based on canonical target sites (Grimson *et al.*, 2007), which are then ranked by weightage. Highest weight is given for an 8mer complementarity, followed by 7mer-m8 (perfect match of positions 2 to 8 of a mature miRNA) and 7mer-1A (exact match of positions 2 to 7 of mature miRNA, followed by an A). The number of mRNA targets are given, followed by numbers of conserved (CS) and poorly-conserved (PCS) binding sites. The total of binding sites often exceeds number of targets, as the candidate miRNA may bind to more than a single site on a single mRNA. Top-three ranked genes for each of the prime CytSh and CytEl-miRNAs are listed, along with gene ontology and expressional traits, in Table 2. Primer pairs designed to specifically (and redundantly, where there are transcript variants and isoforms) target their mRNA in qPCR are listed in Supplementary Table S3. MiRNAs with potential to elicit the 3′UTR of *CTPsynIsoC* were also sought through TargetScanFly ORF. Two of the four prime candidate miRNAs i.e. miR-975 and miR-932 were found to have possible binding sites on the gene’s mRNA, albeit at low probabilities (Supplementary Table S4).

**Table 2:**
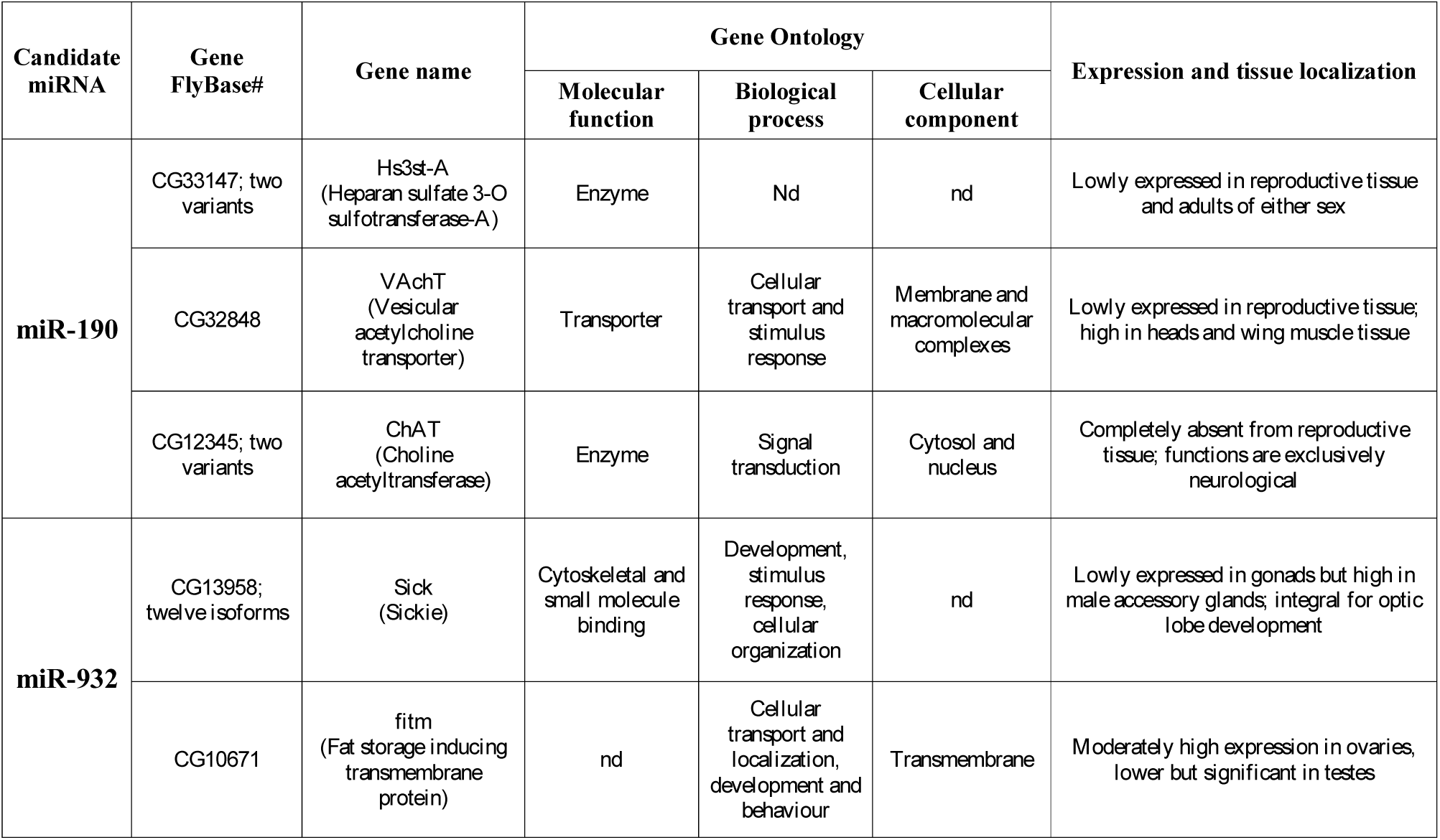
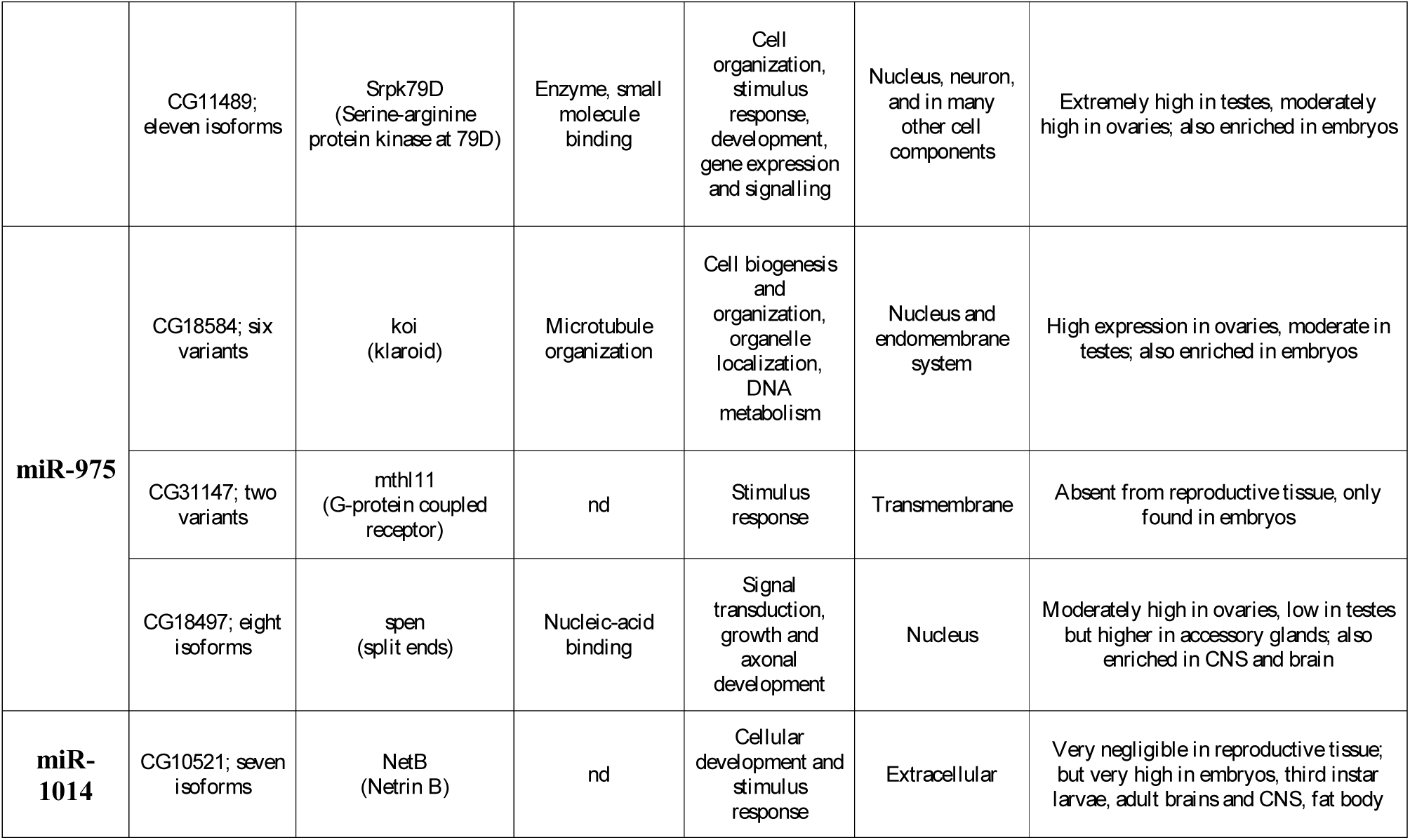
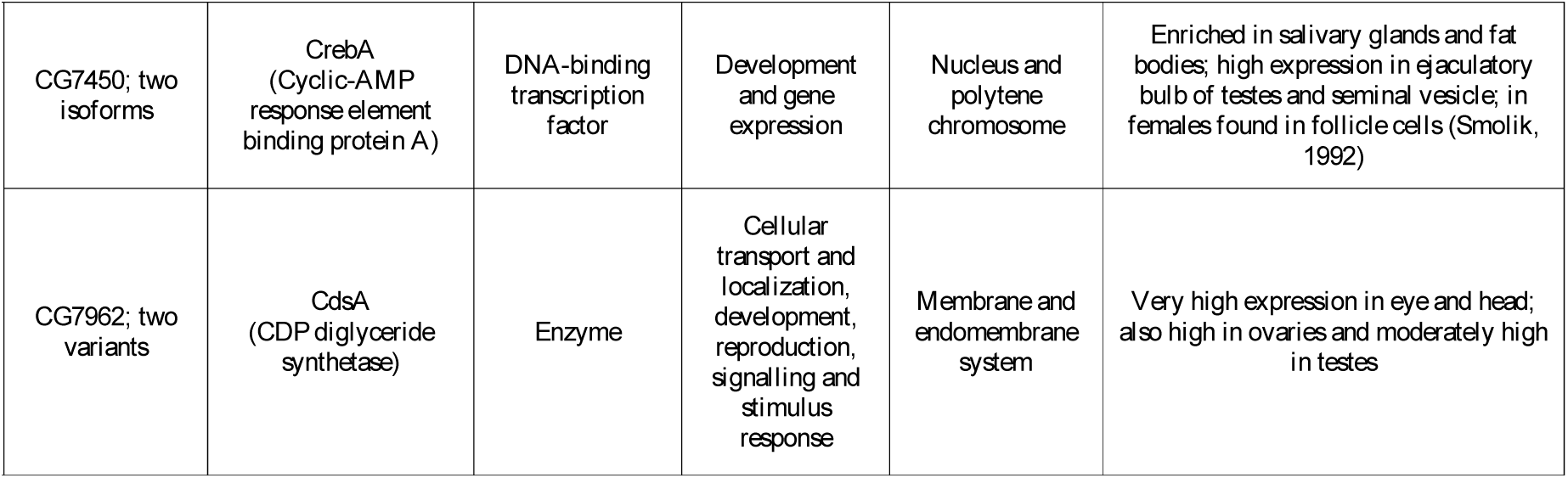
Target predictions of prime candidate miRNAs. ‘nd’ represents not-described. All information is viewable on FlyBase.org through accession with FlyBase#

### 3.6 qPCR analysis against predicted *in silico* targets

In order to determine the target of the miRNAs, qPCR were carried out against the top-three annotated predicted targets specific to each candidate as a means for validation. *CTPsynIsoC* was included as well, to further ascertain whether the overexpression of any of the candidates would affect the gene directly. Results are shown in Figure 9. Controls were either cells transfected with empty pAc-backbone (top) or *Act5c-GAL4>Oregon-R* ovaries (bottom).

**Figure 9:**
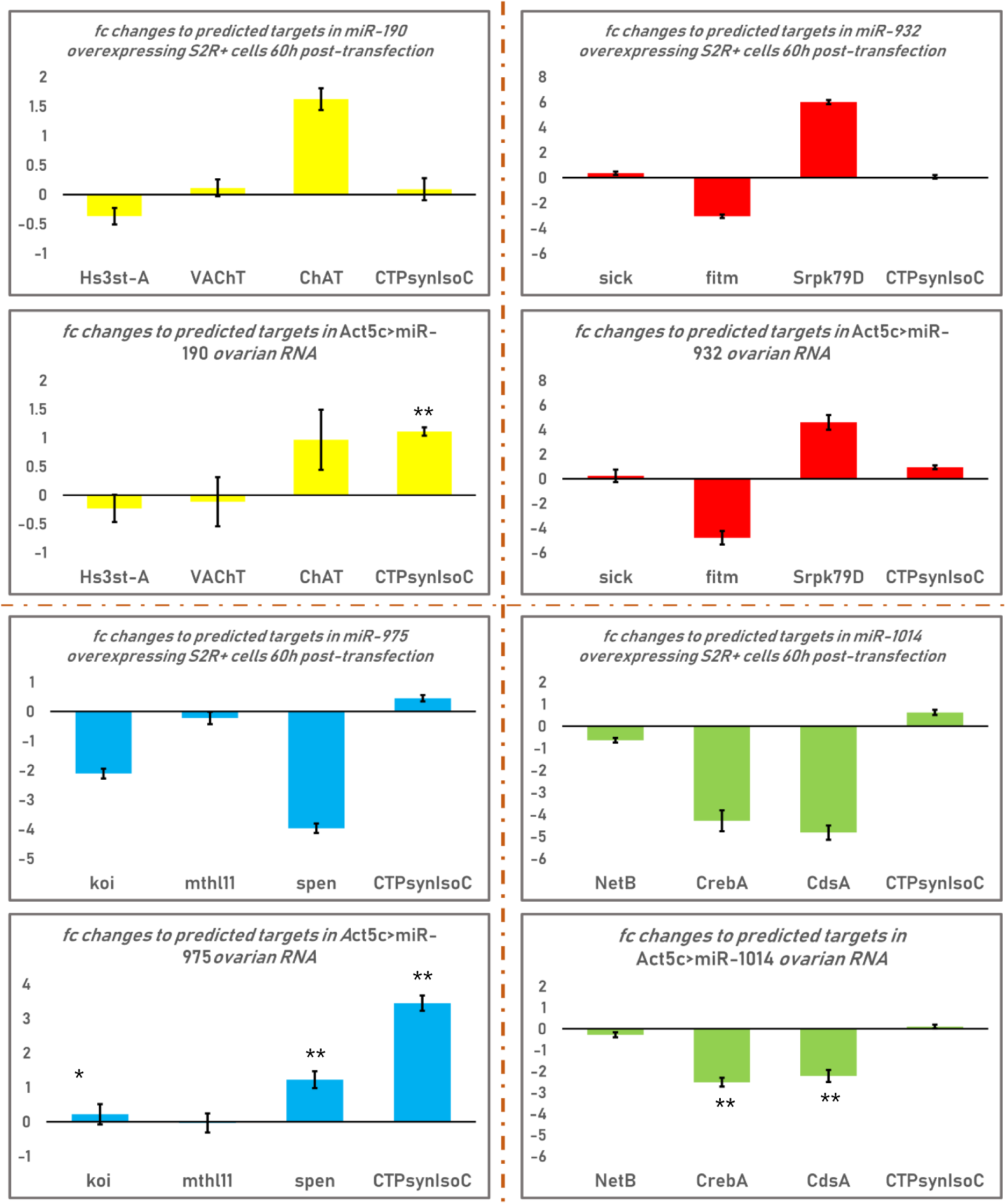
qPCR outcomes of predicted targets in miR-overexpressing cells and miR-overexpressing ovaries. *CTPsynIsoC* quantification is also included. ‘**’ indicates a significant change in expression change to its counterparts e.g. spen in miR-975-overexpressing S2 cells (top) vs spen in miR-975-overexpressing ovaries (bottom). Only the ovarian graph of each miRNAs will be annotated if relevant.

Expressional outliers were apparent right away. For instance, *ChAT* was found to be upregulated in miR-190 overexpressing cells. The same was observed in ovarian tissue, accompanied by significantly greater levels of *CTPsynIsoC*. *Srpk79D* levels also surged in both miR-932-overexpressing cells and ovaries, where its downturn was expected. Otherwise, the expression profile of the four genes under assessment mirrored each other for this CytSh-miRNA, regardless of tissue-type. The same cannot be said for miR-1014-overexpressing tissues. Although all three of its predicted targets were downregulated, reductions to both *CrebA* and *CdsA* in ovarian tissue were considerably less prominent than they were in cell culture. Nevertheless, no differences were more so stark than those conferred by miR-975 in cells as opposed to ovaries: where *koi* was downregulated in cells, it was unchanged in ovaries; where *spen* was downregulated in cells, it was upregulated in ovaries. *CTPsynIsoC* levels were also much higher in *Act5c-GAL4>UAS-miR975* ovaries, with an fc of 3.45 to the meagre 0.45 in cells. These dissimilarities convey that, at least for some miRNAs, differences in the *in vitro* vs *in vivo* microenvironment do indeed play a role in the effects they assert upon gene expression.

## 4.0 Discussion

### 4.1 MiRNA overexpression in follicle cells contributes to non-cell autonomous cytoophidia regulation

Throughout the first stage of screening using follicle cell drivers (FCD), we consistently observed the peculiarity that eventhough GAL4 was only expected to be activated in follicle-cells, nurse cell macro-cytoophidia often appeared affected by miRNA-overexpression. This non-autonomous nature of macro-cytoophidia regulation was also observed to a certain extent within driver-specific controls. Of the three types (FCDs, *nos* and *Act5c*) utilized in this study, only *nos-GAL4>Oregon-R* did not show visibly truncated cytoophidia. We were able to take advantage of this outcome in shortlisting candidate miRNAs; however, its true significance lies in novel aspects of soma-to-germ cells communication it may have shown, specifically between that of follicle and nurse cells of the egg chamber (FNCC). The importance of intercellular communication during embryogenesis is well-understood (Mahowald and Hardy, 1985). The oocyte is dependent on the systematic deposition of maternal factors from nurse cells throughout embryonic development. In the meantime, encapsulating follicle cells provide the oocyte the nutrients it requires for growth. Both examples of “material dumping” occur through specialized gap junction from follicle cells, or ring canals from nurse cells (Cabej, 2012). Comparatively, however, little is known of how FNCC takes place between follicle and nurse cells – if such events happen at all.

Overexpression of candidate miRNAs in follicle cells were consistently apparent within nurse cells. This suggests that FNCC does indeed take place. Two observations especially supported this claim: (A) excessive levels of CytEl-miRNAs (i.e. miR-975 and miR-1014) in follicle cells reversed shortened cytoophidia in *C323a*-driven follicle cells; and (B) despite the proven ineffectiveness of UASt-miRNA constructs in germline cells, effects of miRNA overexpression by FCDs were nonetheless capable of inducing morphological changes to macro-cytoophidia of nurse cells. However, FNCC does not appear to be “two-way”. This was demonstrated through the overexpression of CytSh-miRNA miR-932, when driven by the germline specific driver *nos-GAL4*. Whereas cytoophidia in nurse cells were accordingly truncated, the structures remained of average length in surrounding follicle-cells. Together, these outcomes show that whilst nurse cell cytoophidia could be non-autonomously regulated, their formation in follicle cells are autonomous and unaffected by external stimuli.

Many questions were raised, in which the most pressing one is the component(s) transferred during FNCC. In plants, the plasticity of cellular products transfers such as miRNAs and proteins between neighbouring cells is well-documented (Marin-Gonzalez and Suarez-Lopez, 2012; Zambryski, 2004). Intercellular trafficking occurs through plasmodesmata, channels which are formed through the connective tissue connecting the plant cells (Brunkard, Runkel, and Zambryski, 2015). On the other hand, channel-mediated transfer of macromolecules rarely occur in animal systems. Phospholipid vesicles instead are often used to transfer RNAs and proteins to a nearby cell (Jose, 2015). Furthermore, though follicle cells are physically connected to outer nurse cells in the *Drosophila* egg chamber, ring canals are not known to form in between these two types of cells (Airoldi *et al.*, 2011). Nonetheless, evidence of molecule transfer from follicle to nurse cells do exist. The bacteria *Wolbacchia*, for example, can be transmitted from the somatic to germline stem cell in early-staged egg chambers of various *Drosophila* species. Unfortunately, how the process actually occurs is yet to be explained (Toomey *et al.*, 2013).

Such passive mechanisms of cell-to-cell communication (CCC) are restricted by molecule-size. We therefore hypothesize two scenarios to have most likely caused the non-cell autonomous regulation of nurse cell cytoophidia observed here. One is due to the transfer of excessive candidate miRNAs from follicle to nurse cells, where they go on to affect the same mRNA targets, thereby resulting in the same cytoophidia phenotypes. The second scenario involves CTP molecules themselves. It is known that certain nucleotides are paramount in CCC as signaling molecules (Chen, Levy, and Lightman, 1995; Gründling and Lee, 2016), where much of their function is related to homeostasis maintenance and coordinating metabolic processes (Mediero and Cronstein, 2013; Meshkini, 2014). Moreover, metabolomics-centric studies have shown that the fine nucleotide molecule is much more frequently transported through ring canals and gap junctions than larger cellular components (Pitts and Simms, 1977; Subak-Sharpe, Burk, and Pitts, 1969). CTPsyn enzyme is allosterically regulated by CTP; we therefore reason that as miRNA overexpression upregulate or downregulate CTP level in follicle cells, the passive movement of these molecules across these two cell populations affects CTPsyn activity and eventually, inducing or reducing cytoophidia formation. Our future work will entail nucleotide-movement tracking and quantification assays to assess the validity of this hypothesis, alongside the utilization of fly lines with mutations impairing the proper formation of ring canals, as well as any other types of cell-to-cell channels which may apply.

### 4.2 CTPsyn is not directly regulated by the prime candidate miRNAs

TargetScanFly-mediated target prediction analysis upon the 3′UTR of *CTPsynIsoC* assigned a total of just five potential miRNAs to the region, based on seed complementarity (Schnall-Levin *et al.*, 2010). Regardless, two of the prime candidates i.e. miR-975 and miR-932 were each predicted to have a direct binding site on *CTPsynIsoC*, though neither was conserved beyond the Sophophora subgenus. However, several outcomes of *in vitro* assays conducted within this study has led us to conclude that in reality, neither of these miRNAs may actually do so.

To ascertain availability of true *CTPsynIsoC* 3′UTR binding sites for any of the candidates, a single-miRNA overexpression construct was doubly-transfected alongside a plasmid carrying an *eGFP* gene with the 282-bp 3′UTR sequence of *CTPsynIsoC* attached to its end. Significant eGFP signal reduction compared to controls indicates that a miRNA might indeed have a compatible seed sequence along this region, and therefore could be a direct regulator of *CTPsynIsoC*. The results of this assay demonstrated that miR-975 and miR-1014, and not miR-932, are the ones that are most likely to regulate *CTPsynIsoC*. Its signal reductions were however minute, and *CTPsynIsoC* levels in all miRX-ov cell cultures were not affected. These clues suggest that even if any of these miRNAs were to able regulate the *CTPsynIsoC* gene *in vivo*, effects will be modest and non-significant.

Subsequent qPCR analysis of miR-975-overexpressing ovarian tissues quickly debunked its case as a canonical regulator of *CTPsynIsoC*. Its expression levels not only saw an upturn under such conditions, but the upregulation was significantly drastic. We are hereby more inclined to believe that though *CTPsyn* and cytoophidia formation truly do respond to miRNA-overexpression events, the gene itself is not directly regulated by the candidates under evaluation here. The naturally low expression levels of both miR-975 and miR-932 – as the two most likely to target *CTPsynIsoC* – only provides more support to this assumption. As a matter of fact, if this is to be viewed in lieu to its poor targeting by known D. melanogaster miRNA families, it is not a stretch to say that *CTPsyn* may not be implicitly bound by any miRNAs at all.

Overall, the stringent algorithm of TargetScanFly listed only the potential interactions between five miRNAs and specific sequences within the 3′UTR region of *CTPsynIsoC* to be of any merit. It is important to note that in this case, only the canonical aspects of miRNA-based regulation are considered. For one, the traditionally accepted view has always been that miRNA action would often negatively affect the levels of its mRNA targets. However, this has been challenged, as several miRNA species are found to induce, rather than reduce genetic expression (Catalanotto, Cogoni, and Zardo, 2016; Ørom, Nielsen, and Lund, 2008; Place *et al.*, 2008). Additionally, emerging evidence over the past years has dictated that these small RNAs may assert its regulatory functions by binding to intronic, coding regions, and 5’UTR targets (Lytle, Yario, and Steitz, 2007; Qian *et al.*, 2016). *D. melanogaster* was found to contain these outlying binding sites too (Schnall-Levin *et al.*, 2010). Consequently, with the whole open-reading frame of *CTPsynIsoC* included for target prediction, thirteen miRNAs were identified as potential regulators of the gene. In the future, exploring these routes of *in silico* target prediction as well may be in the best of interests, so that no avenues are left unaddressed.

### 4.3 Influence of fat metabolism pathways in cytoophidia formation

The connections between cytoophidium and fat metabolism become clearer as we go through *in silico* target prediction alongside qPCR validation. This should be expected, given the role of CTP as the seed molecule of phospholipids (Kennedy and Weiss, 1956). Presented outcomes have led us to identify miR-932 and miR-1014 as the two candidates with the most obvious connections to fat metabolism pathways. Based on the expressional fluctuations of their potential targets, we believe that such demonstrated relationships may provide new insights in cytoophidia formation and regulation. These hypothesized pathways are represented by diagrams provided here as Supplementary Figures S2 and S3.

MiR-932 overexpression induces a severe reduction in *fitm* mRNA levels in both ovarian tissue and cell culture. The *D. melanogaster fat-storage inducing transmembrane protein* gene (*dme-fitm*) is homologue to the human *FIT2* gene, in which the protein can be found along the endoplasmic reticulum (ER) membrane (Kadereit *et al.*, 2008). FIT2 acts by partitioning triglyceride (TAG) fatty acids from the ER and packing them into lipid-rich organelles known as lipid droplets (LD). This process occurs when there is a high level of fat within the cells (Guo *et al.*, 2009). Overexpression of *FIT2* increases LD numbers, whilst *FIT2*-knockdown reduces the accumulation of LDs rapidly. On the other hand, miR-1014 overexpression reduces *CdsA* levels. CdsA enzyme is crucial in the initiation of phospholipid synthesis, in which it catalyses the condensation of phosphatidic acid and CTP into the starting molecule i.e. cytidine diphosphate diacylglycerol (CDP-DAG) within phospholipid production (Liu *et al.*, 2014). We hypothesized that cytoophidia elongation may have occurred due to the depletion of CdsA from miR-1014 regulation: as CTP molecules are accumulating, they act on CTPsyn via a negative feedback mechanism (Aronow and Ullman, 1987). CTPsyn thus become inactivated and tether into filaments to avoid degradation, producing the elongated phenotype as observed Nevertheless, what is most crucial here is the effect these events have on the diminishment of phospholipid metabolism, and what that ultimately means for other cellular processes.

For most organisms including *D. melanogaster*, carbohydrates often act as the primary source of energy. However, upon a prolonged starvation period, TAGs will be used in its stead (Owen *et al.*, 1998). To maintain internal TAG level, the plasma membrane breaks off into phospholipid-containing lysosomes within the cytosol, which are promptly rerouted and stored within LDs. These events explain why, in starved cells, not only are LDs larger, but greater numbers of them are found (Rambold, Cohen, and Lippincott-Schwartz, 2015).

It is known that whilst CTPsyn forms the bulk of its cytoophidia, they are not the sole component of the filament (Liu, 2011). The structure was discovered by the anti-Cup antibody, not anti-CTPsyn, indicating there is at least one non-CTPsyn constituent of the cytoophidia (Liu, 2010). Detectable gaps are also found on the macro-cytoophida in the nurse cells, showing its non-homogenous property.

This is where starvation and subsequent fat metabolism may bear links to cytoophidia formation. Under miR-1014 overexpression, the downregulation of *CdsA* eventually depletes ovarian cells of their phospholipid reserves. This creates an environment mimicking starvation which, as aforementioned, culminates in the depletion of phospholipids to compensate for the loss of other fat resources. A previous study has found that elongated cytoophidia were formed in the brains of starved larvae, and at greater frequencies (Aughey and Liu, 2015). True enough, miR-1014-overexpressing egg chambers hold much longer macro-cytoophidia, with the occasional increment in numbers of micro-cytoophidia observed in follicle cells. The same study later showed that once larvae has been refed, cytoophidia disassembling occurred quickly. It can be said that this is mirrored in the truncation of cytoophidia under miR-932 excess: *Act5c-GAL4>UAS-miR932* tissues see much reduced FITM protein levels, thus mimicking conditions of both high-fat food consumption and FIT2-knockdown (Miranda *et al.*, 2014).

*CTPsynIsoC* levels in both *Act5c-GAL4>UAS-miR932* and *Act5c-GAL4>UAS-miR1014* flies were already revealed to be undisturbed. This indicates that CTPsyn molecule availability is not the determining factor to modifications to cytoophidia length in either cases, but are artefacts of the polymerization and tethering processes. As explained beforehand, starvation and fat utilization leads to change in LD level is the most likely cause to a parallel change in cytoophidia behavior. Starvation dynamics explained above dictate that the factor most likely to be changing in parallel to these opposing displays of cytoophidia behaviour are LDs. Apart from lipid storage, these membrane-enclosed organelles have a demonstrated ability to contain and interact with numerous proteins such as histones and membrane-trafficking enzymes (Cermelli *et al.*, 2006). Droplet-recruitment also protects a protein from degradation (Welte, 2007); which is suspected to be one of the main objectives for CTPsyn-filamentation in the first place. Taken together, these are the reasons why we believe LDs to be a literal missing piece of the cytoophidium puzzle. Co-immunoprecipitation, TEM-visualization of tagged-TAG localization patterns, as well as generation and subsequent CTPsyn-centric immunostaining of tissues from fitm-knockout mutants, could be worthwhile endeavours towards figuring out whether this hypothesis holds true.

### 4.4 Summarizing conclusions

Through the systematic utilization of follicle-cell, nurse-cell and ubiquitous drivers, an overexpression-based screening involving over 120 miRNAs has successfully identified a group of candidates which cause either the lengthening or truncation of cytoophidia in *dme-*egg chambers. Both miR-975 and miR-1014 are prime cytoophidia-elongating candidates. Phenotyping revealed that despite this shared trait, they go on to confer differential effects on egg chamber survival: whereas miR-975 overexpression doubles typical nurse cell numbers in a small number of egg chambers, miR-1014 significantly increased incidence of apoptosis instead. Conversely, prime candidates miR-190 and miR-932 only appeared to cause cytoophidia-shortening. As morphological changes to these filamentous structures appear little dependent on *CTPsyn* levels themselves, these observations together signify that rather than cytoophidia-dissociation, CTPsyn polymerization events may be a more useful indicator of major changes to the cellular ‘normal’. Subsequent *in vitro* assays further revealed very low probabilities of any of the four prime candidates eliciting the cytoophidia-forming isoform of *CTPsyn* i.e. *CTPsynIsoC*, suggesting the participation of intermediary components. Whilst *in silico* prediction followed by qPCR validation of targets showed feasible *CTPsyn* connections for neither miR-190 nor miR-975, they uncovered the possibility of a deep-seated relationship between fat metabolism and cytoophidia formation, to miRNAs 932 and 1014. All-in-all, this study has successfully demonstrated the involvement of a small group of miRNAs in CTPsyn regulation, although it is highly likely that their roles in these cases are indirectly conferred.

## Acknowledgement

We would like to thank all our collaborators and colleagues for the discussion and the work conducted in this lab. This study was funded by the Fundamental Research Grant Scheme (203/PBIOLOGI/6711457). N.D. is funded by Yayasan Khazanah.

